# Description of *Alternaria kierstenii* sp. nov. from invasive garlic mustard highlights agricultural biosecurity risks of foreign weeds as pathogen reservoirs

**DOI:** 10.64898/2026.11.02.763701

**Authors:** Cole Sawyer, Jami L. Thomas, Glen Groben, Michael R. Fulcher, Matthew A. Tancos

**Author notes:** The author(s) declare no conflict of interest.

## Abstract

Invasive weeds can increase agricultural biosecurity risks by serving as reservoirs for emerging or undercharacterized plant pathogens. Garlic mustard (*Alliaria petiolata*), an invasive brassicaceous weed in North America, occurs near agricultural landscapes and is susceptible to multiple crop pathogens. A previously undescribed *Alternaria* species was isolated from a diseased garlic mustard population in Maryland, USA, prompting investigation of its taxonomy, origins, and host range. *De novo* genome sequencing produced a near-chromosome scale assembly of 35.8 Mb comprising 17 contigs. Comparative genomic analyses with other crucifer-infecting *Alternaria* species revealed expanded repertoires of predicted CAZymes and peptidases, as well as unique effectors and biosynthetic gene clusters. Phylogenetic and comparative analyses supported recognition of this fungus as *Alternaria kierstenii* sp. nov., representing a distinct lineage within section *Sonchi.* Host range assays demonstrated that *A. kierstenii* was highly aggressive on diverse hosts in the Brassicaceae, including vegetable, ornamental, cover crop, and invasive species. The disease caused by this new pathogen manifested as foliar and stem lesions, chlorosis, necrosis, leaf death, and, in several hosts, rapid plant mortality. Amplicon sequence data from a nationwide garlic mustard microbiome study detected closely related taxa only in the plants from the eastern states of Maryland, Pennsylvania, and New York, suggesting a geographically restricted population consistent with expectations for a potentially emerging pathogen. The description of this phylogenetically unique, broad host range pathogen demonstrates the potential for widespread invasive weeds to harbor pathogens and pathogen genomic diversity with agricultural relevance, underscoring the hidden biosecurity risk of understudied invasive weeds.

## INTRODUCTION

Invasive species cost the global economy US$1.288 trillion, threatening agriculture, forestry, ecosystem diversity, infrastructure, and public health (Diagne et al., 2021). Biological invasions are a major avenue for emergent diseases as non-native species are introduced into new regions, altering ecosystems and introducing novel microbiota and pathogens (Zhang et al., 2022). Animal invasions are a leading cause of emerging zoonotic diseases as previously allopatric species are permitted to interact in time and space at higher frequencies or durations, increasing the probability of microbial movement, pathogen evolution, and animal-to-human host jumps that threaten public health (Zhang et al., 2022). Similarly, the movement of non-native plant species not only results in the introduction of agricultural and ecological pests (i.e., weeds), but also the introduction or accumulation of novel plant-associated microbes, viruses, and pathogens, with unknown agricultural biosecurity risks (Tancos et al., 2024). Non-native weeds grown at monoculture levels adjacent to taxonomically related crops increase the risks of pathogen movement, spillover, and emergence events due to the introduction and/or selection of more aggressive strains or entirely new phytopathogenic species (Dubrow et al., 2022; Fulcher et al., 2024, 2026; Linde et al., 2016; Tancos et al., 2024). Predicting and identifying these biosecurity risks is challenging due to an inadequate amount of research investigating invasive plant disease ecology and the multitude of uncharacterized microbes and viruses that non-native weeds can introduce or harbor (Keskitalo et al., 1998).

The mustard plant family (Brassicaceae) contains numerous non-native invasive weed species that critically impact US agriculture and inhabit large portions of the US. Prominent examples include hoary alyssum (*Berteroa incana* (L.) DC.), hoary cress (*Lepidium draba* L.), perennial pepperweed (*Lepidium latifolium* L.), hairy whitetop (*Lepidium appelianum* Al-Shehbaz), dyer’s woad (*Isatis tinctoria* L.), Sahara mustard (*Coincya tournefortii* (Gouan) Alcaraz, T.E.Díaz, Rivas Mart. & Sánchez-Gómez), yellow rocket (*Barbarea vulgaris* W.T.Aiton), and garlic mustard (*Alliaria petiolata* (M.Bieb.) Cavara & Grande) (DiTomaso, 2000; Durka et al., 2005; Francis & Warwick, 2008; Gaskin et al., 2018; Hurka et al., 2003; MacDonald & Cavers, 1991; Rejmanek et al., 1991; Rodriguez et al., 2024; Rollins, 1981; Warwick & Francis, 2006). While these harmful members of the Brassicaceae are included across many state and federal invasive weed management programs, the family also includes high-value vegetables like broccoli, cabbage, cauliflower, and kale. These crops have a combined value of US$2.4 billion and an average value of US$12,254 per acre, more than 15 times the per-acre value of grain corn, the highest-valued US commodity crop (USDA, 2025a, 2025b). Additionally, the brassicaceous oilseed crops canola, rapeseed, and mustard seed are valued at US$1.0 billion and are harvested from over 2.9 million acres across the US (USDA, 2025a, 2025b). The extensive cultivation of Brassicaceae crops across the US, combined with the co-occurrence of diverse non-native brassicaceous weeds and their associated pathogens, may increase the risk of new crop disease epidemics and pathogen movement between cultivated and non-cultivated hosts.

The cruciferous biennial garlic mustard has been studied extensively in a plant invasion biology context (Callaway et al., 2008; Rodgers et al., 2022) and, similarly, may represent a useful model system for investigating pathogen emergence in invasive plant populations. The utility of the system for this purpose is derived from its (i) widespread presence and ecological damage, (ii) ability to naturally harbor diverse crop pathogens, (iii) overlapping distribution with brassicaceous crops, and (iv) target for biological control programs (Durka et al., 2005; Katovich et al., 2022; Tancos et al., 2024; Tancos & Frederick, 2020). European garlic mustard populations were introduced into North America in the 19^th^ century and have colonized temperate forests and agricultural hedgerows in over 36 states and the District of Columbia (Bennett et al., 2020; Durka et al., 2005; Marshall, 1989; Rodgers et al., 2022). Garlic mustard rapidly displaces and outcompetes native or beneficial plant species due to its allelopathic properties, lack of herbivores, and high fecundity (Alabi et al., 2021; Bennett et al., 2020; Durka et al., 2005; Meekins & McCarthy, 1999). While few North American herbivores consume garlic mustard, plant pathogens have exploited this widespread weed, evidenced by natural infections with the crop pathogens *Xanthomonas campestris*, *Cucumber mosaic virus*, *Turnip mosaic virus*, *Turnip vein-clearing virus*, *White clover mosaic virus*, and *Erysiphe cruciferarum* (Boehm & Nameth, 2000; Fulcher & Owen-Smith, 2022; Gaafar et al., 2019; B. Lockhart et al., 2014; B. E. Lockhart, 2012; Tancos & Frederick, 2020; Zhao et al., 2016). In addition to natural field infections, greenhouse host range studies have also demonstrated garlic mustard’s susceptibility to multiple crop-pathogenic *Xanthomonas* isolates (Tancos et al., 2024). Recently, a novel pathogenic *Alternaria* species was identified on a heavily diseased garlic mustard population in the US state of Maryland and evaluated as a potential biological control agent for garlic mustard (Tancos et al., 2026). The origin of the previously undescribed *Alternaria* species remains unknown but could be from (i) pathogen acquisition during invasion, (ii) pathogen co-introduction with garlic mustard, or (iii) *de novo* evolution (Tancos et al., 2026).

*Alternaria* is a fungal genus in the Pleosporales with over 368 accepted species separated into 29 sections (Li et al., 2023). *Alternaria* taxa can be found worldwide, which comprise a cosmopolitan group of species that act as saprophytes, endophytes, or pathogens of animals and plants (Vinogradova et al., 2026). Phytopathogenic *Alternaria* species typically produce blighted symptoms that include necrotic leaf spots, chlorosis, defoliation, and rot, leading to reduced crop marketability and yield reduction (Thomma, 2003). Broad host ranges, toxin production capacity, and agricultural impact make *Alternaria* species notorious plant pathogens (Thomma, 2003). Agriculturally important *Alternaria* species include, but are not limited to, *A. brassicicola* (hosts: Brassicaceae), *A. citri* (hosts: Rutaceae), *A. dauci* (hosts: Apiaceae), *A. solani* (hosts: Solanaceae, Cucurbitaceae), *A. triticina* (hosts: Poaceae), and *A. zinniae* (hosts: Asteraceae) (Akimitsu et al., 2003; Gannibal et al., 2022; Koike et al., 2017; Nowicki et al., 2012; Schmey et al., 2024; Thomma, 2003; Tralamazza et al., 2018). At least ten *Alternaria* species have been documented as pathogens on hosts in the family Brassicaceae, yet the *Alternaria* sp. nov. from garlic mustard belongs to a new monotypic sister lineage in section *Sonchi* and is most closely related to *A. triangularis*, a host-specific pathogen characterized in Japan on *Bupleurum* (Apiaceae) (Nishikawa & Nakashima, 2020; Tancos et al., 2026). While garlic mustard is not considered an agronomic weed that directly competes with field crops, its widespread distribution, proximity to crops via hedgerows, and susceptibility to diverse crop pathogens all increase the agricultural biosecurity risks of this ‘woodland’ weed.

The evolutionary origin of this fungus remains unresolved. It may have been acquired from a pathogen already present in North America, co-introduced with garlic mustard, or evolved *de novo* following invasion (Tancos et al., 2026). We hypothesized that *Alternaria* sp. nov. FDWSRU 22-162 represents a genetically distinct *Alternaria* lineage with genomic features relevant to Brassicaceae pathogenicity. To investigate its evolutionary relationships and pathogenic potential, we (i) sequenced and annotated the FDWSRU 22-162 genome, (ii) assessed its evolutionary relationships and pathogenicity-associated gene diversity, (iii) characterized its experimental host range, and (iv) examined the geographic occurrence of closely related taxa using amplicon data from garlic mustard microbiomes. These analyses provide evidence relevant to the proposed origin hypotheses, characterize the agricultural host range of this novel pathogen, and clarify the potential agricultural biosecurity risks associated with invasive garlic mustard.

## MATERIALS AND METHODS

### Fungal strain and culture conditions

*Alternaria kierstenii* sp. nov. FDWSRU 22-162 was maintained on potato dextrose agar (PDA, Neogen, Lansing, MI, US) and incubated under ambient laboratory temperature and lighting (∼20-24°C), as previously reported (Tancos et al., 2026). While *in vitro* sporulation was demonstrated, it was sparse and of an insufficient quantity for host range studies. Therefore, a macerated mycelial suspension was used for inoculations (Tancos et al., 2026). To prepare mycelial suspensions, two agar plugs from an actively growing culture were suspended in 2 mL sterile distilled water and homogenized (TissueLyser II, Qiagen, Germantown, MD, US). A 200 µL aliquot of the homogenized suspension was then used to inoculate 200 mL of potato dextrose broth (PDB, Neogen, Lansing, MI, US). Liquid cultures were incubated under ambient laboratory conditions on an Orbital Genie shaker (Scientific Industries, Bohemia, NY, US) at 100 rpm. After 7 days, liquid cultures were pooled, and mycelium was collected using vacuum filtration. A 24.5g aliquot of mycelium was added to 400mL sterile distilled water with 0.2% Tween 20 (Sigma-Aldrich, St. Louis, MO, US) and homogenized (IKA® ULTRA-TURAXX® disperser, IKA Works, Inc., Wilmington, NC, US). The suspension was then filtered through a single layer of sterile cheesecloth.

### Sporulation

Artificial sporulation was induced according to methods described in Simmons 2007. *Alternaria kierstenii* sp. nov. FDWSRU 22-162 was point inoculated onto potato carrot agar (PCA) (Simmons, 2007). Petri plates were incubated unsealed at 22°C under a daily fluorescent light/dark cycle of 8/16 hr. After 7 days, conidia were gently scraped from the agar surface and mounted on a slide in lactic acid. Features were observed and measured under a compound light microscope (Olympus BX41, Olympus Corporation, Center Valley, PA, US) at 400X magnification. Length and width of conidia, conidiophores, and conidial beaks (n=25) were measured (NIS-Elements, Version 5.30.02, NIKON Instruments, Inc. Melville, NY, US) from two independent plates and averaged.

### Host range trials

Experimental plants used in pathogenicity tests are listed in Supplementary Table 1. Garlic mustard seed require a cold, moist stratification period to germinate. Therefore, field-collected seeds from healthy plants were incubated on moist filter paper within a black plastic bag and refrigerated (4°C) until seeds germinated (approx. 3 months) (Tancos et al. 2024). Seedlings were then transplanted into 4-inch diameter square black form pots (T.O. Plastics, Inc., Clearwater, MN, US) containing a commercial soilless mix (Sunshine Mix #1, Sun Gro, Agawam, MA, US) amended with a controlled release fertilizer per manufacturer specifications (Osmocote 14-14-14, The Scotts Company, Marysville, OH, US). Following garlic mustard germination, all other experimental plants were direct sown into prepared 4-inch diameter pots.

One-month old plants, with at least three fully developed true leaves, (*n =* 5) were spray inoculated with macerated mycelium suspended in sterile water and 0.2% Tween 20 (10^3^ CFUs mL^-1^) until runoff. Negative control plants (*n =* 5) were sprayed with sterile water with 0.2% Tween 20. Treated seedlings were placed in an artificial dew production chamber for 48-hours at 100% RH and 20°C and then maintained under greenhouse conditions (20-25°C) with natural light in a completely randomized layout. Disease incidence was recorded seven days post inoculation. All experiments were independently repeated at least twice.

### Analysis of infection data

Data were analyzed in Rstudio v.2026.01.1+403 using R v.4.5.1. A generalized linear model of infection severity, defined here as the number of diseased leaves divided by the total number of leaves per plant across both trials, was produced using the binomial logit function (R Core Team, 2025). Post-hoc comparisons of severity between different hosts were then performed using the emmeans package (version 2.0.4) and converted into compact letter display using the included multcomp::cld() function (Lenth & Piaskowski, 2017).

### Genome sequencing and assembly

High molecular weight genomic DNA was extracted from fresh *A. kierstenii* sp. nov. FDWSRU 22-162 mycelium being grown in liquid PDB using the protocol developed by Penouilh-Suzette et al. 2020 (Penouilh-Suzette et al., 2020; Tancos & Fulcher, 2024). DNA quality and quantity was assessed using a Qubit fluorometer (Thermo Fisher Scientific, MA, USA) and a FEMTO Pulse electrophoresis system (Agilent, CA, USA). Extracted DNA was sent to the Delaware Biotechnology Institute at the University of Delaware DNA Sequencing & Genotyping Center for PacBio HiFi (Pacific Biosciences, CA, USA) library preparation and long-read sequencing using PacBio Revio. Assembly was completed using 1,335,281 reads with a total length of 18,909,407,915 and N50/N90 = 14749/10083. A kmer analysis of the raw reads was performed with GenomeScope 2.0 with kmer length 21, and max kmer coverage set to 10000. The genome was assembled *de novo* using Flye v 2.9.4 with a kmer length of 17 and minimum overlap requirement of 10,000 bp (Kolmogorov et al., 2019). An initial read coverage of 100x was specified, which resulted in the use of only reads ≥ 19,293 bp in length. Mitochondrial contigs were removed from the final assembly for separate analysis. BUSCO v 5.7.1 was run on the nuclear genome using the euk_genome_min prediction mode and the Dothideomycete_odb10 database to estimate completeness. The mitogenome was annotated with MFannot (Lang et al., 2023), MITOS2 (Fiedler et al., 2023), GESeq (Tillich et al., 2017), RNAweasel (Lang et al., 2007), and manual blast searches of open reading frames identified by ORF-FINDER (Rombel et al., 2002). Consensus annotations were curated using Geneious Prime v 2025.0.1 (GraphPad Software, Boston).

### Genome annotation and comparative genomics

Assembled fungal genomes were downloaded from NCBI in March 2025 and annotated using funannotate v.1.9.0 (Palmer & Stajich, 2020) with default parameters. Functional analysis of protein coding regions was performed using Interproscan v.5.78-109.0 (Jones et al., 2014). Carbohydrate-active enzymes (CAZymes) and peptidases were predicted via blastp searches against dbCAN and MEROPS databases respectively with a cut-off value of 60% identity and 60% coverage (Rawlings et al., 2018; Zheng et al., 2023). Secondary metabolite gene clusters were annotated using antiSMASH v.8 (Blin et al., 2025). For putative effector analysis, secreted proteins were predicted using SignalP v.6. Proteins containing transmembrane helices and GPI anchors, as predicted by TMHMM v.2 and PredGPI v.1, were removed from the analysis (Pierleoni et al., 2008; Teufel et al., 2022). The final subset of proteins was run through EffectorP v.3 (Sperschneider & Dodds, 2022). Analysis of macrosynteny between all five fungal genomes was performed using MCScan through the JCVI toolkit v.1.5.3 (Tang et al., 2024). Genome repeat content was determined using RepeatModeler v.2.0.4 (Flynn et al., 2020).

### Annotation of starship transposons

Starfish v.1.1.0 was used to predict putative tyrosine recombinases in all five fungal genomes and locate potential starship transposons (Gluck-Thaler & Vogan, 2024a). Predicted starships were manually searched for the well-characterized starship-associated proteins annotated as ankyrin repeat protein, heterokaryon incompatibility protein, patatin-like phosphatases NB-ARC domain proteins, myb/SANT domain protein or NLR domain protein (Gluck-Thaler et al., 2022; O’Donnell et al., 2025). Synteny analysis was performed by aligning the predicted starship regions using minimap2 v.2.24-r1122 and visualized with gggenomes v.1.1.3. To collect proteins found within the tandem repeat region across the fungal cohort, protein sequences were collected and used as input for tblastn via Geneious prime v.21.0.7+6-LTS with a threshold of 60% identity and 60% coverage (Camacho et al., 2009). Regions were collected, aligned and visualized in the same manner as the putative starships.

### Garlic mustard microbiome analyses

The potential geographic distribution of *Alternaria kierstenii* was preliminarily determined by comparing partial ITS2 sequences with amplicons generated during an independent study of garlic mustard microbiome diversity (NCBI GenBank SRA: PRJNA1474556 (Fulcher et al. in review). Briefly, this separate study characterized microbiomes from 250 asymptomatic garlic mustard samples collected in twelve states, including California, Colorado, Maryland, Minnesota, New York, North Carolina, Oregon, Pennsylvania, Rhode Island, Vermont, Washington, and Wisconsin. In Maryland, intensive sampling at three locations focused on garlic mustard compartments, which comprised bulk soils, rhizospheres, roots, lower leaves, stems, upper leaves, and seed pods. At the other 11 states, only upper leaves were sampled. At each site, replicate samples were collected from five plants found within a 15 m^2^ area. Metagenomic DNA was extracted from each sample using a commercial kit, and ITS2 (5.8SR: TCGATGAAGAACGCAGCG) and ITS4:TCCTCCGCTTATTGATATGC) libraries were prepared and sequenced at the University of Minnesota Genomics Center on an Illumina NextSeq 2000 with a P2 flow cell outputting 300 bp paired-end reads.

Amplicon sequencing data were processed using qiime2 version amplicon-2024.10 to generate amplicon sequencing variants (ASVs), cluster operational taxonomic units (OTUs) at 97% similarity, and assign taxonomy based on the UNITE reference database (Abarenkov et al., 2024; Bokulich et al., 2018; Bolyen et al., 2019; Callahan et al., 2016; Rognes et al., 2016). ASVs assigned to the genus *Alternaria* and closely related genera—*Stemphylium*, *Bipolaris*, *Curvularia*—were considered in the present study, and only those *Alternaria* ASVs with a frequency of 100 reads in one or more samples were retained for analysis (Mainali et al., 2017). These ASVs and the corresponding partial ITS2 sequence from *A. kierstenii* FDWSRU 22-162 were used in a phylogenetic analysis. Sequences were aligned with MAFFT version 7.490, and the resulting alignment was analyzed with IQ-TREE v.2.2.3 to generate maximum likelihood trees with automatic model selection and 1,000 ultrafast bootstraps (Hoang et al., 2018; Katoh & Standley, 2013; Minh et al., 2020). A consensus phylogenetic tree using a 70% support threshold was created with GeneiousPrime v.2023.0.1 (Biomatters Inc., Boston MA USA). The tree was rooted to the ASV assigned to the *Curvularia* genus, and *Alternaria* ASVs were labelled with their corresponding 97% OTU.

## RESULTS

### *Alternaria kierstenii* has a broad host range and is highly aggressive on Brassicaceae

Diverse brassicaceous species, representing vegetables, ornamentals, a cover crop, garlic mustard, and the solanaceous crops tomato and pepper, were evaluated for susceptibility to *A. kierstenii* sp. nov. FDWSRU 22-162. Disease symptoms, including foliar and stem lesions and leaf death, were documented across all brassicaceous species (Table 1, Figure 1). Symptoms progressed rapidly with plant death occurring on garlic mustard, cabbage, stock, and wallflower (Figure 1). No disease symptoms were documented on any control plants. Leaf breakage was commonly observed on field mustard following inoculation due to the leaf’s large surface area, dew treatment, and abundance of stem lesions. In contrast, leaf breakage did not occur on control field mustard replicates. The solanaceous crops, tomato and pepper, were similarly investigated, resulting in disease symptoms on tomato, but not pepper (Table 1, Figure 1). Disease symptoms were less severe on tomato, compared to brassicaceous hosts, primarily producing foliar lesions and a few incidences of stem lesions. *Alternaria kierstenii* sp. nov. FDWSRU 22-162 was successfully reisolated from all tested leaf samples from diseased cabbage, garlic mustard, and tomato.

**Figure 1.**
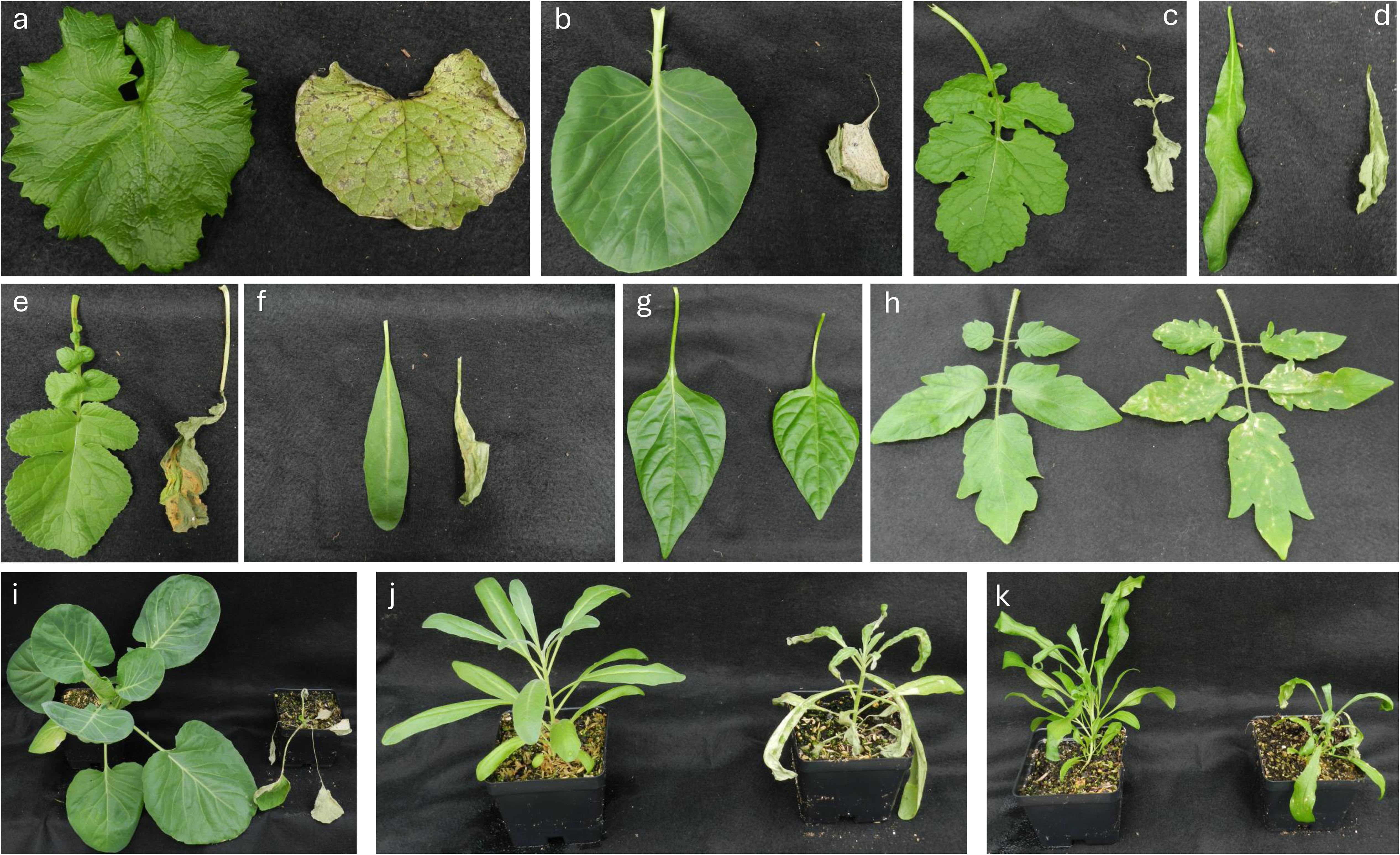
Host range of *Alternaria kierstenii* sp. nov. Sub-panels a-h are representative leaf samples from mock inoculated (left) and inoculated (right) plants, while sub-panels i-k are representative whole plants. a) Garlic mustard, b) cabbage, c) field mustard, d) wallflower, e) radish, f) common stock, g) pepper, h) tomato, i) cabbage, j) common stock, k) wallflower.

**Table 1.** Disease incidence of *A. kierstenii* sp. nov. across two independent trials.

| **Species** | **Diseased Plants** | **Stem Lesions** | **Diseased Leaves** | **Severity (%)** | **Significance<sup>†</sup>** |
|---|---|---|---|---|---|
| Pepper | 0/10 | 0/10 | 0/69 | 0.00 | - |
| Tomato | 9/10 | 2/10 | 35/67 | 52.24 | b |
| Stock | 10/10 | 10/10 | 90/107 | 84.11 | a |
| Radish | 10/10 | 10/10 | 57/65 | 87.69 | a |
| Garlic | 10/10 | 10/10 | 42/47 | 89.36 | a |
| Wallflower | 10/10 | 10/10 | 102/114 | 89.47 | a |
| Cabbage | 10/10 | 9/10 | 63/67 | 94.03 | a |
| Field Mustard | 10/10 | 10/10 | -<sup>*</sup> | -<sup>*</sup> | <sup>-</sup> |

<sup>\*</sup>Significant stem damage prevented proper classification of leaf samples
<sup>†</sup> Significance determined using generalized linear model as outlined in methods

### Genome of the emergent brassicaceous plant pathogen *Alternaria kierstenii* sp. nov

GenomeScope predicted a genome of 37,677,239 - 37,689,447 bp with a repeat length of 7,789,595 - 7,792,119 bp. The nuclear genome was predicted to be haploid, and one non-nuclear (mitochondrial) genome was expected. However, the final assembled mitogenome was resolved into two circular contigs (Supplementary Figure 1). One larger contig of 48,022 bp with 29.6 % GC content (497x coverage), and a smaller contig of 13,886 bp with 29.4% GC content (26,646x coverage). The extremely high coverage recorded for the smaller contig suggests that either a true two-chromosome mitochondrial genome is present with a chromosome copy disparity or extreme sequencing bias resulted in a discontiguous assembly of a single chromosome. Removal of mitochondrial contigs prior to further analysis produced a nuclear genome of 35,790,988 bp with 50.9% GC content (575x average contig coverage).

The *A. kierstenii* sp. nov. FDWSRU 22-162 nuclear genome was assembled into 17 contigs. The assembly is near-complete, with single gene copies detected for 3,752 of 3,786 BUSCO groups (99.1%), no duplicate groups detected, five fragments found, and 29 groups missing. Based on the presence of telomeric repeats on contig ends (TTAGGG_5_), seven complete chromosomes were resolved, and three putative partial chromosomes were detected with greater than 1 Mb length and telomeric repeats on one terminus each (Figure 2a). Seven smaller contigs (13,439 to 84,007 bp) were assembled, three of which also contained telomeric repeats on one terminus each (Figure 2b). The remaining four contigs may represent discontiguous portions of core chromosomes but could also represent dispensable or accessory chromosomes.

**Figure 2.**
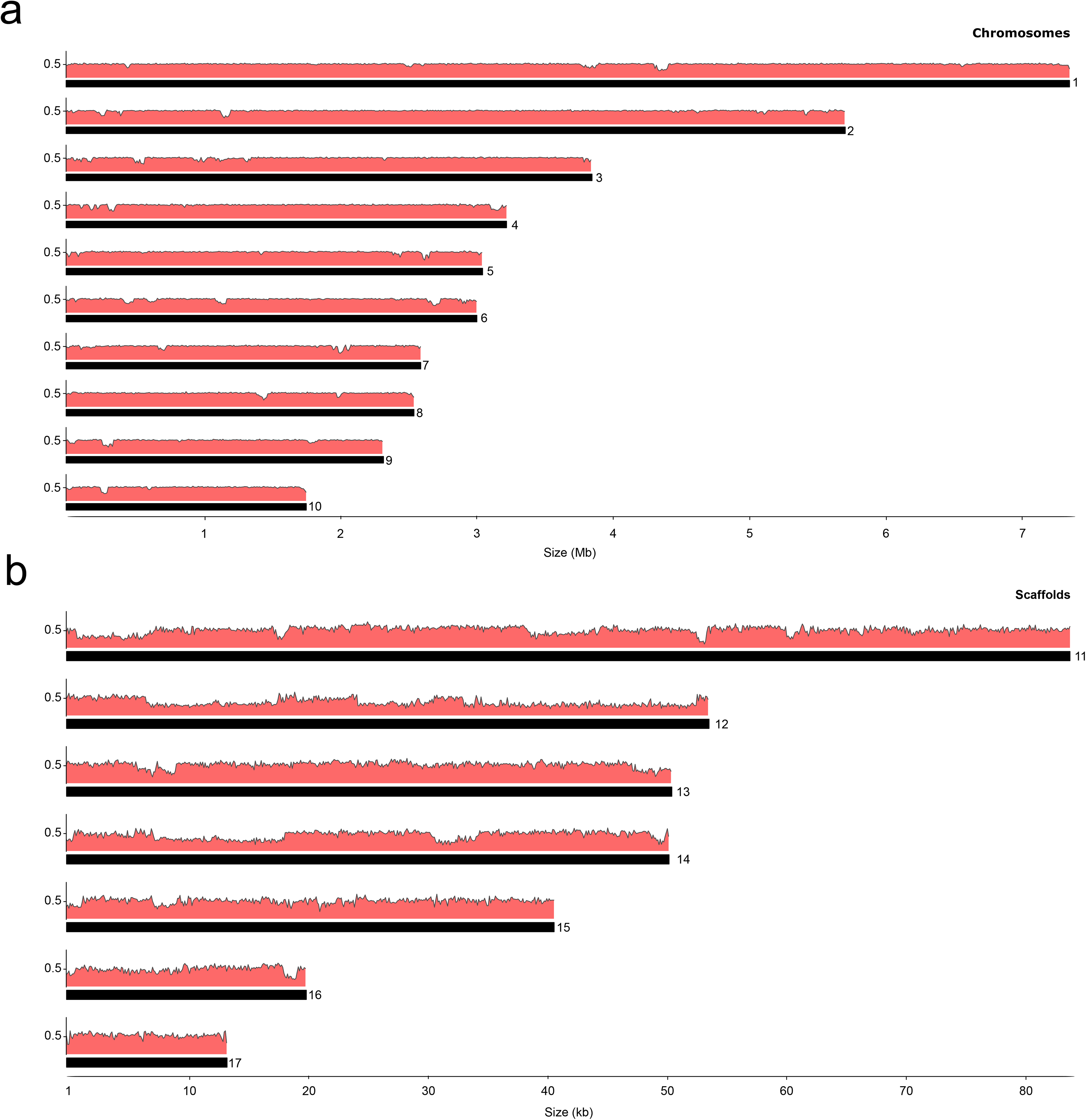
Genome of *Alternaria kierstenii* sp. nov. Ideogram of chromosomes (a) and scaffolds (b) of the *A. kierstenii* genome. Each bar represents the approximate length of each chromosome/scaffold. The GC content is displayed above the corresponding chromosome/scaffold with a mid-line placed at 50% GC content. Chromosome sizes are measured in megabases (Mb) while scaffold sizes are measured in kilobases (kb)

### *Alternaria kierstenii* sp. nov. possesses a large genome with an expanded number of putative effectors compared to other *Brassica*-infecting *Alternaria* species

We compared the chromosomes of our novel *Alternaria* species to those of known *Brassica* crop pathogens (*A. brassicae* J3 and *A. brassicicola* Abra43), as well as two strains of the model species *A. alternata*. Strain PN1 was isolated alongside *A. brassicae* J3 and found to be avirulent in the absence of *A. brassicae* while strain Z7 is a pathogen of tangerine (Gai et al., 2021). Using this cohort, we sought to highlight the characteristics which are unique to brassicaceous plant infecting species and locate unique genetic factors carried by *Alternaria kierstenii* sp. nov. In terms of size, the genome of *A. kierstenii* sp. nov. FDWSRU 22-162 was the largest (35.8 mb), similar in size to *A. brassicae* J3 (34.1 mb). Genome alignments found synteny across the ten core chromosomes common within *Alternaria* species, but no homology was found between the *A. kierstenii* sp. nov. FDWSRU 22-162 genome assembly and known accessory chromosomes of *A. brassicae* J3 and *A. alternata* Z6 (Figure 3a). Although the shorter contigs from *A. kierstenii* sp. nov. FDWSRU 22-162 showed no homology to the other *Alternaria* genomes, these may represent distinct and novel accessory chromosomes (Figure 3a). Analysis of repeat content found that all three pathogens associated with the Brassicaceae, *A. kierstenii* sp. nov. FDWSRU 22-162, *A. brassicae* J3 and *A. brassicicola* Abra43, had a greater number of predicted transposable elements relative to the *A. alternata* strains (Supplementary Figure 2). A large inversion was noted on chromosome 3.

**Figure 3.**
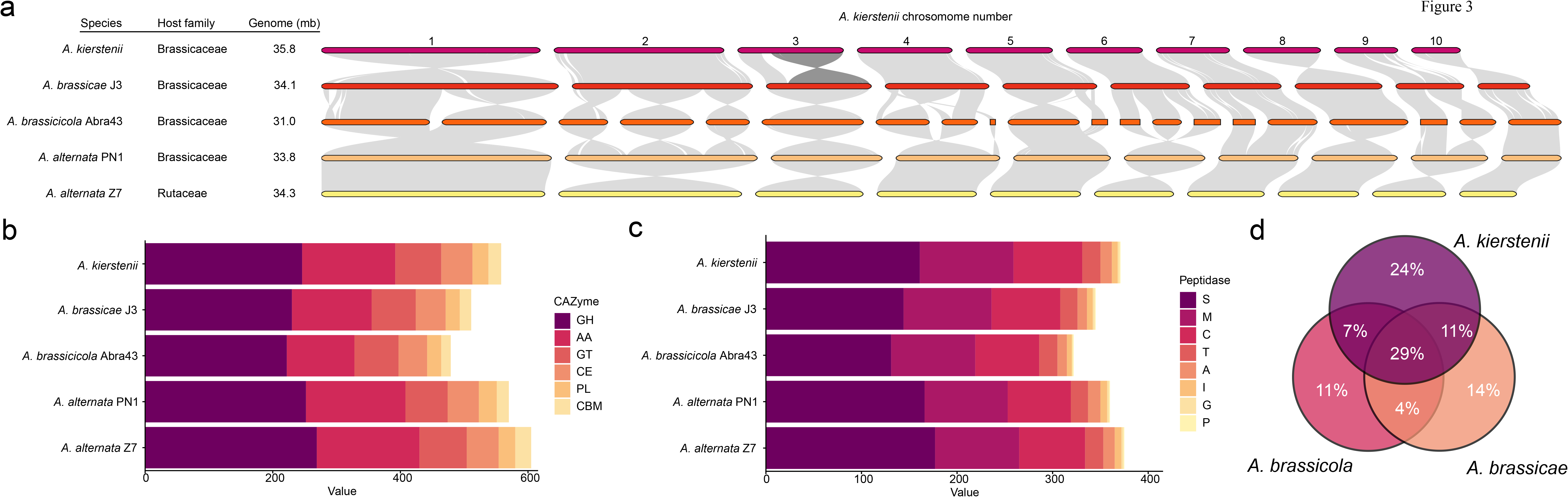
Genome comparisons of *Alternaria kierstenii* sp. nov. to other brassicaceous *Alternaria* species. a) Comparisons of plant host source, genome size and synteny. Grey bands represent homology between syntenic blocks of chromosomes/scaffolds. A large inversion of chromosome 3 of *A. kierstenii* is colored dark grey. Stacked bar charts of CAZymes (b) and peptidases (c) for all *Alternaria* species. d) Venn diagram comparing effector proteins of brassicaceous infecting *Alternaria* species.

Comparisons of gene repertoire reveal that *A. kierstenii* sp. nov. FDWSRU 22-162 had an expanded number of both CAZymes (Figure 3b) and peptidases (Figure 3c) when compared to the other brassicaceous plant infecting fungi. These increases were dispersed throughout the different sub-classes of CAZymes and peptidases, with no single group showing specific evidence of enrichment. Of the 246 effectors predicted from the *A. kierstenii* sp. nov. FDWSRU 22-162 genome, 100 were shared with both *A. brassicae* J3 and *A. brassicicola* Abra43, while 83 were unique (Figure 3d). Many of these effectors (n = 112) contain no known conserved domains. Of those with annotations, 43 were predicted to encode CAZymes, including 15 AA9, 24 pectin lyase-like proteins and four cutinases. Other putative effectors include two copies of the *Alternaria* major allergen protein 1, two cerato-ulmin type hydrophobins, four chitin deacetylases, two LysM domain effectors and one necrosis inducing protein.

### Secondary metabolite prediction highlights the presence of multiple toxin clusters

Thirty-nine biosynthetic gene clusters (BGCs) were predicted from the *Alternaria kierstenii* sp. nov. FDWSRU 22-162 genome via antismash (Blin et al., 2025). Five were unique among examined *Alternaria* genomes (Figure 4a). Eleven of the clusters predicted in *A. kierstenii* sp. nov. FDWSRU 22-162 were found exclusively in *Brassica*-infecting fungi of our cohort. These include two hybrid clusters composed of more than one backbone enzyme. The first hybrid, cluster 4-5, contained three separate backbone enzymes, two NRPS synthetases and one PKS (Figure 4b). The full cluster was only found in *A. kierstenii* sp. nov. FDWSRU 22-162 and *A. brassicae* J3, with the other species having lost one or two of the backbone proteins. The second BGC, cluster 6-1, was unique in that it contains a dimethylallyl tryptophan synthase (DMAT). The DMAT is paired with a large NRPS backbone and is only found in *A. kierstenii* sp. nov. FDWSRU 22-162 and *A. brassicicola* Abra 43 (Figure 4c).

**Figure 4.**
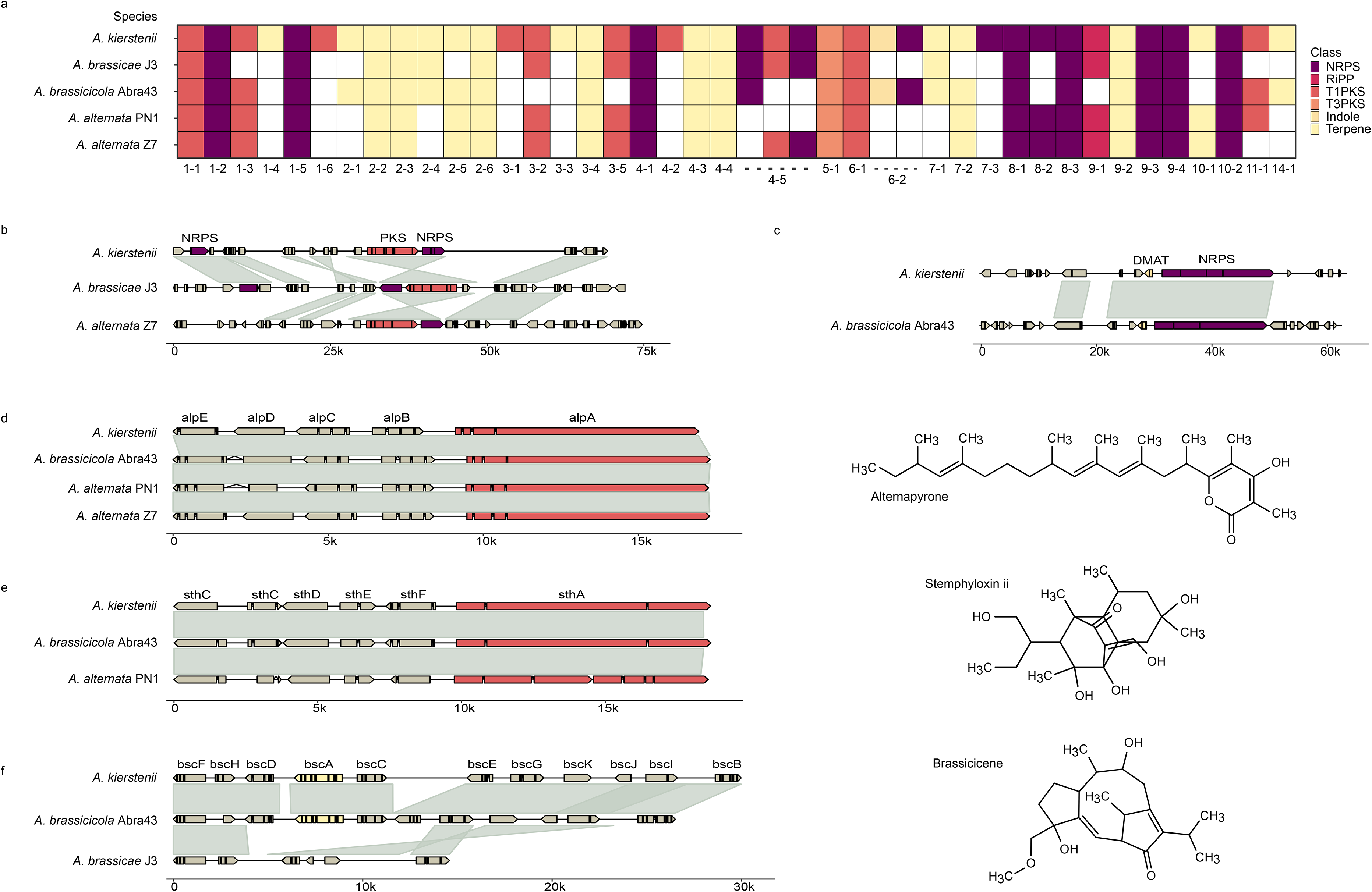
Secondary metabolome of *Alternaria kierstenii* sp. nov. a) Presence absence analysis of the all predicted secondary metabolite gene clusters of *A. kierstenii* sp. nov. across the fungal cohort. Colored blocks represent presence of a given backbone enzyme, with the hue representing the class of molecule synthesized. The number of the cluster in *A. kierstenii* sp. nov. is listed below, hyphens denote hybrid clusters containing more than one backbone enzyme. b-f) Genetic maps of clusters of interest as follows; multi-backbone hybrid cluster 4-5 (b), DMAT containing hybrid cluster (c), alternapyrone cluster (d), stemphyloxin ii cluster (e) and brassicicene cluster (f). Grey ribbons represent homologous syntenic blocks between gene clusters. Predicted molecules synthesized are placed to the right of sub-panels d-f.

Antismash predicted multiple toxin clusters which had been previously found in other plant pathogenic species, specifically clusters 1-3, 11-1 and 14-1 (Figure 4d-f). First, cluster 1-3 was predicted to synthesize the compound alternapyrone, a known phytotoxin (Figure 4d). Using cBlaster, we determined the conservation of this and other clusters in our cohort and found that the alternapyrone cluster is largely conserved but is missing in *A. brassicae* J3 (Figure 4d). Second, Cluster 11-1 shares homology to a cluster from the pathogen *Stemphylium botryosum* which synthesizes the phytotoxin stemphyloxin ii and is found additionally within *A. brassicicola* Abra43 and *A. alternata* PN1 (Figure 4e). Finally, Cluster 14-1 contains all proteins necessary to synthesize the fusicoccane-type terpene brassicicene (Figure 4f), which was previously found to be synthesized by *A. brassicicola* and belongs to a class of known plant phytotoxins. A related cluster in *A. brassicae* J3 is missing the backbone terpene cyclase *bscA* and is likely unable to synthesize the compound.

### A tandem duplication of a Tc-1 mariner transposon within a large starship carries multiple virulence-associated proteins

Examination of an inversion found on chromosome 3 located a large gap in the synteny between *A. kierstenii* sp. nov. FDWSRU 22-162 and *A. brassicae* J3. Analysis of the conserved domains found within this gap identified a DUF3435 tyrosine recombinase, which has been associated with large transposons termed starships. Like integrative and conjugative elements in bacteria, starships are large mobile elements that can carry cargo between fungal genomes. Computational prediction of starship elements in our fungal genome cohort did not locate any starship elements in the *Brassica*-infecting *Alternaria* species but did annotate two starships within each *A. alternata* species (Supplementary Table 2). Starfish did, however, predict that the DUF3435 recombinase of *A. kierstenii* sp. nov. FDWSRU 22-162 was homologous to previously annotated starship captain proteins (E-value 6e-50). A manual search downstream of the captain protein found multiple starship-associated proteins, namely a heterokaryon incompatibility protein, an ankyrin repeat protein, and a myb/SANT domain protein. As myb/SANT proteins have been shown to associate with the end of starship transposons, we categorized the region between the DUF3435 recombinase and the myb/SANT protein as the cargo (Figure 5a) (O’Donnell et al., 2025). Using all proteins of the putative starship as query, we performed a tblastn search against the genomes of our fungal cohort to look for related elements. This analysis returned an element in the genome of *A. brassicae* J3 which carried 10 homologous proteins including the DUF3435 captain and the myb/SANT domain protein (Figure 5a).

**Figure 5.**
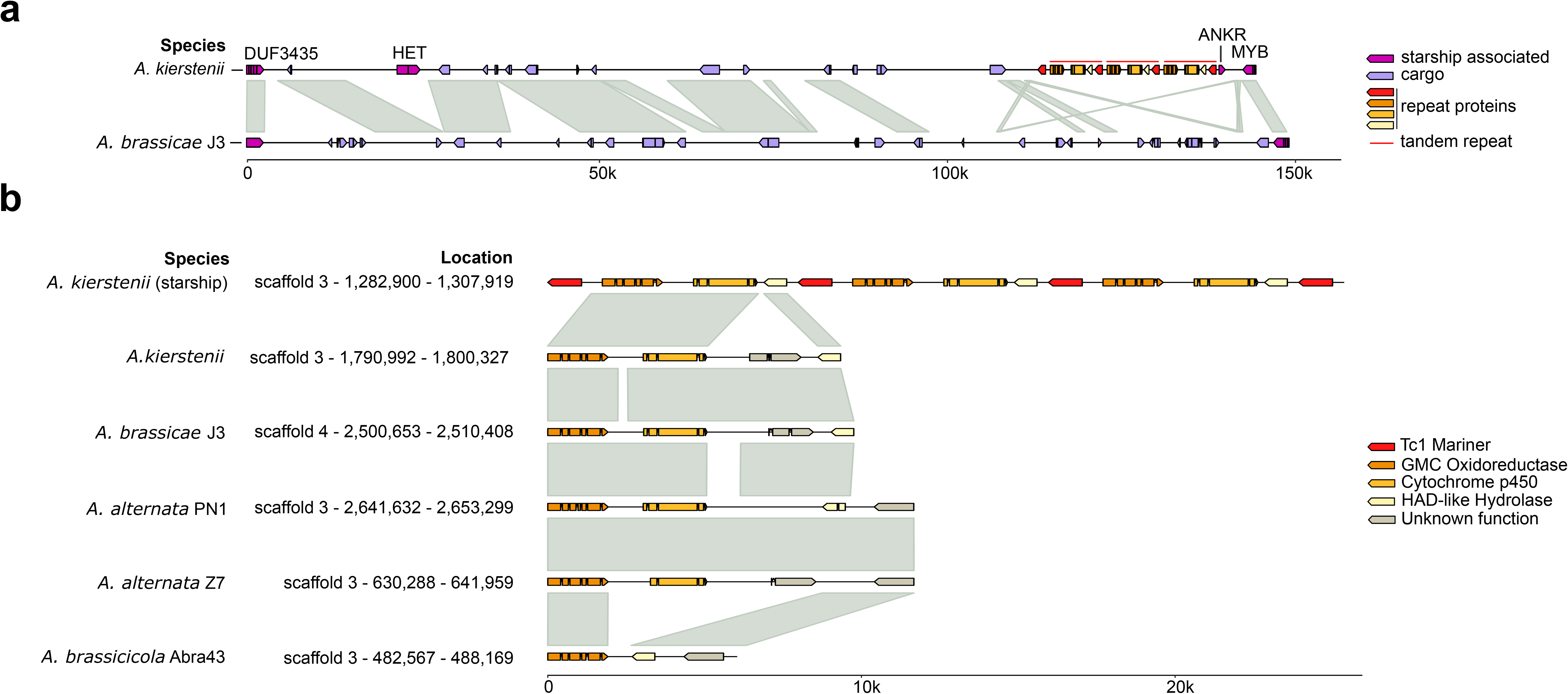
Analysis of a novel starship found in *Alternaria kierstenii* sp. nov. a) Genetic map of the starship found next to the inversion on chromosome 3 of *A. kierstenii* sp. nov. Protein coding sequences previously associated with starship transposons are colored dark purple and are labeled above. Three tandem repeats composed of a Tc1 mariner transposon (red), a GMC Oxidoreductase (dark orange), a cytochrome p450 (light orange) and a HAD-like hydrolase (yellow), are denoted with a red line. b) Genetic maps of the tandem repeats within the starship of *A. kierstenii* sp. nov. compared to non-starship loci in other *Alternaria* species. Coloration follows the same pattern as in sub-panel a and homologous syntenic blocks are represented as grey ribbons. Synteny with the tandem repeats is limited to only the first repeat to aid in readability. Sizes in both panels are measured in kilobases.

Of the 25 proteins denoted as cargo, eight had no known conserved domains, while the remainder contained multiple proteins which may facilitate infection, including a transcription factor, a calcium dependent channel protein, and a cupin. Additionally, we noted three tandem repeats, each consisting of four proteins. Analysis of the conserved domains found that the repeat contained a putative Tc-1 mariner transposon, followed by a glucose-methanol-choline (GMC) oxidoreductase, a cytochrome p450 protein and a haloacid dehalogenase-like hydrolase protein (Figure 5b). Of these, only the GMC oxidoreductase, which itself is an AA3 class CAZyme, contained a signal peptide for export. To determine the origin of these proteins, we then searched for homologs in our fungal cohort. All proteins within the repeat, except for the Tc1-mariner transposase, have homologs in the other *Alternaria* species and at a separate location in *A. kierstenii* sp. nov. (Figure 5b) found in a syntenic group.

To locate similar elements from across the Dothideomycetes, a cblaster analysis was performed. This analysis located multiple putative starship elements; however, many had differing cargoes and/or tyrosine recombinase proteins. For example, the best match was a putative starship from *Pyrenophora tritici-repentis* (Supplementary Dataset 1) with 12 protein homologs but no homology to the recombinase. Additionally, this analysis found that the three repeat element proteins listed previously are commonly found throughout the genus *Alternaria* as a single copy.

### Identification of *A. kierstenii-*like endophytes in asymptomatic garlic mustard populations located across the eastern United States

A separate, nationwide garlic mustard microbiome study presented a unique opportunity to investigate the possible presence and/or prevalence of fungi closely related to *A. kierstenii* sp. nov. FDWSRU 22-162 across the continental United States. In total, 18 unique *Alternaria* ASVs, with at least 100 reads for a given microbiome sample, were identified across 50 of 250 garlic mustard microbiome samples. These 18 unique ASVs collapsed at a 97% similarity threshold to just three *Alternaria* OTUs. Between the three *Alternaria* OTUs, 11 states were represented, including California, Colorado, Maryland, Minnesota, New York, North Carolina, Oregon, Pennsylvania, Rhode Island, Vermont, and Wisconsin. *Alternaria* OTUs 1 and 2 were only identified from leaf material, while OTU 3 was identified in the soil, rhizoplane, leaves, stems, and seed pods of garlic mustard.

*Alternaria* OTU 1 was the rarest and represented only two *Alternaria* ASVs. One of these two *Alternaria* ASVs (973a98c0b36e4e4932de20da1eb23073) was a 100% sequence match (263/263 bp) to *A. kierstenii* sp. nov. FDWSRU 22-162 (Figure 6). Similarly, a BLASTn search in May 2026 using this ASV as a query produced a 100% sequence similarity match to isolates FDWSRU 22-162 and 22-163 with the next closest *Alternaria* sequence at 99% similarity. The *A. kierstenii*-like ASV was recorded in two samples collected from the upper leaves of symptomless garlic mustard populations located in Pennsylvania and New York. In both samples, the *A. kierstenii*-like ASV represented the majority of total *Alternaria* ASVs with over 13,784 reads from the Pennsylvania population and 349 reads from the New York population, representing 81.7% and 86.8% of total *Alternaria* ASVs, respectively. The second *Alternaria* ASV (f43151ebd716fa86794aeea1b65db66b) from OTU 1 had 98.86% sequence identity (260/263 bp) to *A. kierstenii* sp. nov. FDWSRU 22-162 and originated from a Maryland population with 972 reads, representing 83.9% of total *Alternaria* ASVs.

**Figure 6.**
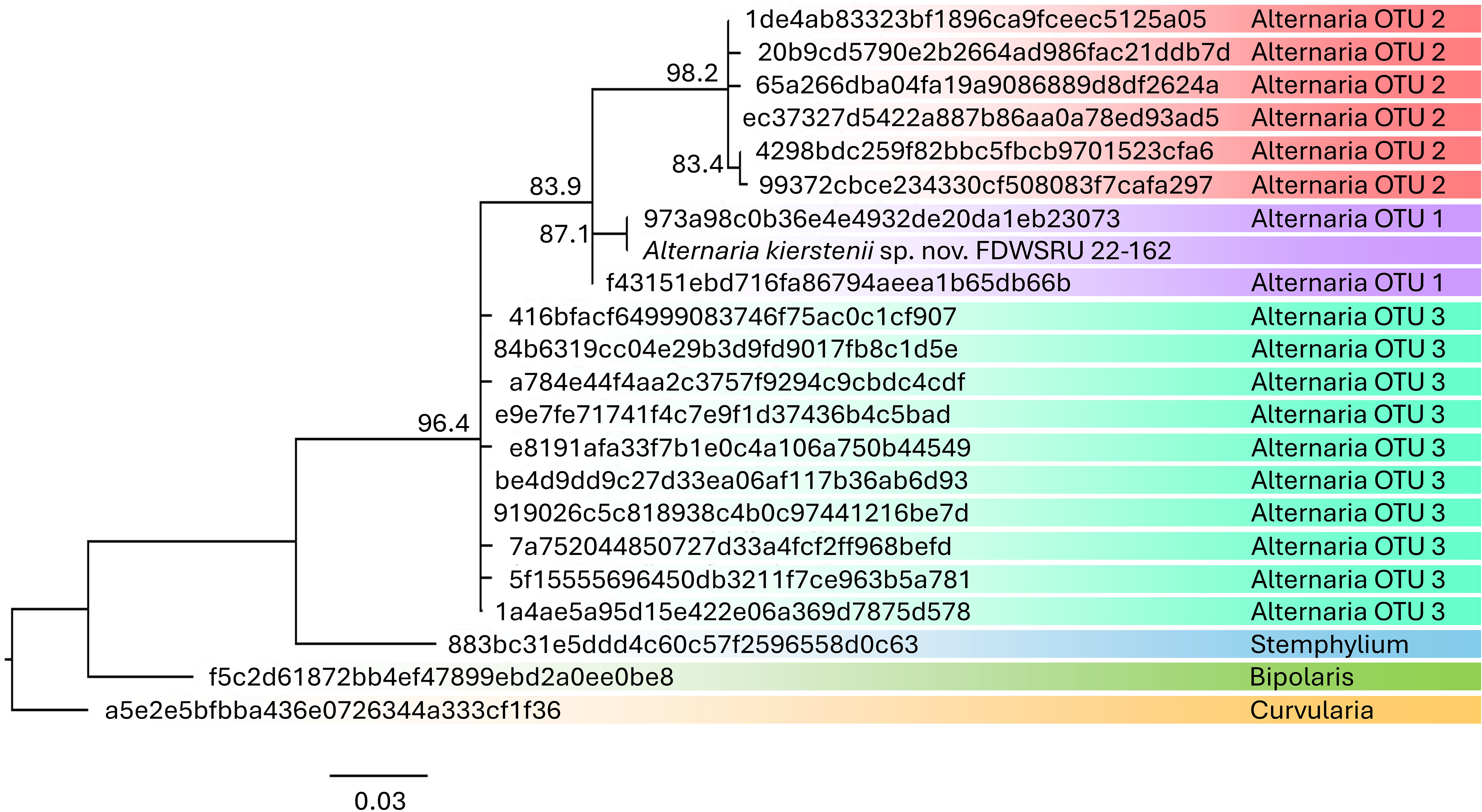
Maximum likelihood phylogenetic tree generated from the internal transcribed spacer (ITS) region 2 from ASVs with at least 100 reads/sample from garlic mustard microbiomes sampled across twelve states across the USA. *Alternaria kierstenii* sp. nov. FDWSRU 22-162 was added for reference. The scale bar represents the number of nucleotide substitutions per site.

In contrast, *Alternaria* ASVs from OTUs 2 and 3 were more geographically widespread across the continental United States. *Alternaria* OTU 3 was the most widespread and isolated from all tissue types and soil. Pairwise sequence identity to *A. kierstenii* FDWSRU 22-162 ranged between 95.49% (254/266 bp) to 96.24% (256/266 bp) for the ten representative ASVs. *Alternaria* OTU 2 was similarly widespread across the continental United States, but in contrast to OTU 3, was limited to garlic mustard leaf tissue. Pairwise sequence identity to *A. kierstenii* sp. nov. FDWSRU 22-162 ranged between 95.09% (252/265 bp) to 95.77% (249/260 bp) for the six representative ASVs.

### Taxonomy

Based on the results from the morphological study conducted here and the previous phylogenetic analysis (Tancos et al., 2026), we described one new species in *Alternaria* Sect. *Sonchi*. The species is described as follows:

*Alternaria kierstenii* M. Tancos, *sp. nov*.

MycoBank: 864287

*Etymology: kierstenii*, in dedication to the corresponding author’s wife, Kiersten A. Tancos.

*Classification:* Pleosporaceae, Pleosporales, Dothideomycetes.

*Description:* Colonies on PDA are slow growing with an average 9.8mm radial growth after seven days at 23℃, white and greyish mycelia growth, irregular form, raised elevation, undulate margins. Hyphae are hyaline to lightly pigmented with acute angle branching and septations; sparse sporulation (Tancos et al. 2026). On PCA, conidiophores, 14.45-123.69µm x 3.61-7.03µm (average 56.65 x 4.96µm), are septate, pigmented pale brown to brown, branched, straight to flexous, wider and darker than vegetative hyphae (Supplementary Figure 3). Conidia are small, 16.76-50.48µm x 6.99-21.06µm (average 32.46 x 11.50µm), muriform, obclavate, pale to dark brown with transverse (1-8) and longitudinal septations (common). Conidia are singular and in chains of (3-)4-8(-10). Lateral branches are present, but minor. Conidium body is constricted near septations. Conidium beak, 3.66-15.84µm x 3.22-6.88µm (average 7.05 x 4.94µm), is medium and blunt to tapered. Secondary conidiophores are short and unbranched.

*Typus:* United States, Mid-Atlantic Region, Maryland, Montgomery County, Silver Spring, Sligo Creek near the Colt Terrace Neighborhood Park on diseased foliage of *Alliaria petiolata* (Brassicaceae), 24 May 2022, *M. Tancos FDWSRU 22-162* (ex-type) (Tancos et al., 2026); USDA Ag Data Commons genome assembly: https://figshare.com/s/5aeee14a5a4b68a7148c. Genbank: ITS PQ375868; actin PQ375864; *Alt a1* PQ375867; ATPase 1 PQ375863; calmodulin PQ375865; *gpd* PQ375866.

*Distribution:* Mid-Atlantic United States.

*Additional specimens examined:* USA, MD, on diseased foliage of *Alliaria petiolata* (Brassicaceae), 24 May 2022, *M. Tancos FDWSRU 22-163*. Genbank: ITS PV231460; actin PV231456; *Alt a1* PV231459; ATPase 1 PV231455; calmodulin PV231457; *gpd* PV231458.

*Notes: Alternaria kierstenii* sp. nov. was isolated from irregular shaped necrotic lesions surrounded by chlorotic halos on the non-native invasive weed *Alliaria petiolata*. An *A. petiolata* infestation of approximately 300 sq ft was severely blighted (100% disease incidence) with severe defoliation and dieback (Tancos et al. 2026). Pathogenicity trials on diverse Brassicaceae and Solanaceae species demonstrated a broad host range, infecting all tested Brassicaceae species and the Solanaceae species tomato (*Solanum lycopersicum*). Genetic analyses place *A. kierstenii* as a monotypic sister lineage to section *Sonchi*, and distinct from closely related *Brassica*-infecting *Alternaria* species (Tancos et al., 2026).

## DISCUSSION

*Alternaria kierstenii* sp. nov. was originally investigated due to its potential as an endemic, host-specific biocontrol agent for garlic mustard management (Tancos et al., 2026); however, following the presently reported host range testing, it became clear that *A. kierstenii* sp. nov. was a poor biocontrol candidate. When evaluated against a representative panel of diverse cultivated hosts, *A. kierstenii* sp. nov. infected all tested Brassicaceae members and a Solanaceae host, tomato. Despite a relatively low inoculum concentration of 10^3^ CFUs mL^-1^, cabbage, stock, and wallflower were highly susceptible, resulting in severe leaf and/or plant death. The rapid onset of leaf and stem necrosis suggests the production of phytotoxins, which are heavily represented across the *Alternaria* taxa (Li et al., 2023; Meena & Samal, 2019). Pepper was the only species tested that showed no susceptibility, whereas tomato was susceptible, but did not develop severe infection like that observed with brassicaceous hosts. *Alternaria* species commonly display broad host ranges (Thomma, 2003). For example, *A. alternata* can infect brassicaceous and solanaceous hosts, but is often considered an opportunistic, weak pathogen and can occur in complexes with more virulent species, such as *A. brassicae, A. brassicicola,* and *A. solani* (Baltzakis et al., 2024; Davis et al., 2026; Rajarammohan, Paritosh, et al., 2019; Rajarammohan, Pental, et al., 2019). In contrast, *A. brassicae, A. brassicicola,* and *A. solani* are predominately associated with their respective solanaceous or brassicaceous hosts (Simmons, 2007). To our knowledge, limited examples exist of *A. brassicicola* or *A. brassicae* infecting tomato (Adama et al., 2021).

Initial attempts at inducing artificial sporulation by *A. kierstenii* were unsuccessful (Tancos et al., 2026), but minor sporulation could be achieved, resulting in the conidial characterization of this small-spored, slow-growing species. The closest taxonomically related species, *A. triangularis,* is also a slow-growing, sparse sporulator, producing short to moderately long chains of small conidia, like *A. kierstenii* sp. nov. (Nishikawa & Nakashima, 2020; Tancos et al., 2026). The only other *Brassica*-infecting *Alternaria* species within section *Sonchi* is the slow-growing *A. brassicae*, which produces chains of large conidia (Nishikawa & Nakashima, 2020; Rajarammohan, Pental, et al., 2019). There are no available reference genomes for *A. triangularis*, and few complete genomes are available for any *Alternaria* species associated with the Brassicaceae, limiting the investigation of genomic factors underlying host-microbe interactions. Therefore, the high-quality genome assembly produced by this study will be useful specifically during investigation of *A. kierstenii* sp. nov. evolutionary origins and more broadly for understanding molecular factors critical to Brassicaceae plant infections that might be conserved across *Alternaria* species.

Long-read sequencing generated a genome for *A. kierstenii* sp. nov. containing 10 putative chromosomes, consistent with the genome architecture of other *Alternaria* species, as well as seven additional scaffolds (Ushijima & Yamamoto, 2019). These scaffolds failed to align to either core or accessory chromosomes of *A. brassicae* J3 or *A. alternata* Z7 suggesting that these elements may represent i) novel accessory chromosome(s) or ii) accessory regions of the core chromosomes which failed to assemble. The size of the scaffolds, ranging from 13 to 84 kbp, the large number of predicted transposons, and the presence of telomeric sequences corresponding to the number of missing telomeres on the core chromosomes, suggests the latter for three of the shorter contigs (Bertazzoni et al., 2018). Accessory regions are critical to many plant pathogens and often encode toxins associated with pathogenicity such as the T-toxin PKS of *Cochliobolus heterostrophus* (Kodama et al., 1999). The exact location of these scaffolds in the genome as well as their association with virulence is currently under study.

Comparative analyses between *A. kierstenii* sp. nov. and the *Alternaria* species of our cohort demonstrated that *A. kierstenii* sp. nov. had the largest genome, largely owing to expansive repeat content which is a commonality in *Brassica*-infecting species. Similarly, both the CAZyme and the peptidase repertoire of *A. kierstenii* sp. nov. were enlarged when compared to *Brassica*-infecting species suggesting that *A. kierstenii* sp. nov. is likely able to process more plant derived compounds during infection as seen in other plant pathogens (Baroncelli et al., 2016; Brunner et al., 2013). Additionally, *A. kierstenii* sp. nov. contained the largest effector pool of the *Brassica*-infecting species, with 83 unique to this species when compared to our cohort. While the relationship between virulence factors and host range can be difficult to parse due to the presence of gene-for-gene interactions, our computational data and greenhouse experiments suggest that *A. kierstenii* sp. nov. may have a broader host range than other *Brassica*-infecting species, though further study of side-by-side infection processes is required.

As *Alternaria* species are prolific producers of host-specific toxins, we generated a secondary metabolome using antismash (Hussain et al., 2025; Tsuge et al., 2013). This analysis found that the BGCs of *A. kierstenii* sp. nov. encode multiple previously characterized toxins, specifically alternapyrone, stemphyloxin ii and brassicicene (Ichihara, Oikawa, Hayashi, et al., 1983; H. Li et al., 2018; Phakeovilay et al., 2022; Tazawa et al., 2018). Alternapyrone belongs to the class of α-pyrones, which contain many known mycotoxins such as citreoviridin and asteltoxin, while alternapyrone was shown to have antigermination effects on wheat seed (El-Sawy et al., 2023; H. Li et al., 2018; Lin et al., 2016). Stemphyloxin ii was previously noted as a phytotoxin in the plant pathogen *Stemphylium botryosum* and is structurally related to betaenone compounds which induce chlorosis on host leaves (Ichihara, Oikawa, Hashimoto, et al., 1983; Ichihara, Oikawa, Hayashi, et al., 1983; Ugai et al., 2015). Lastly, brassicicene, a molecule first described in *A. brassicicola* belongs to the family of fusicoccane diterpenoids (J.-Y. Liu et al., 2022; Minami et al., 2009). These molecules include the namesake compound fusicoccin, a phytotoxin which induces wilting in plant species, and are structurally similar to ophiobolin, another phytotoxic compound (Evidente, 2024; J.-Y. Liu et al., 2022). The activity and phytotoxicity of these compounds in *A. kiersteni*i sp. nov. are currently uncharacterized, however, given the literature surrounding these metabolites, we hypothesize that these metabolites are likely to contribute to the virulence of *A kierstenii* sp. nov. on multiple hosts. Our analysis also identified clusters 4-5 and 6-2 which each contain multiple backbone enyzmes. Of particular interest is the DMAT containing cluster 6-2, as these proteins synthesize various bioactive alkaloid compounds (Metzger, 2009). The metabolites synthesized by these clusters are currently unknown, however, it is possible that the metabolites produced will similarly impact fungal pathogenicity on brassicaceous hosts. Future research should focus on determining which clusters are active during infection and their putative roles.

Comparing the genomes of these *Alternaria* species led to the discovery of a large genomic inversion in *A. kierstenii* sp. nov. Within this region, we identified a tyrosine recombinase (DUF3435) which had previously been associated with large transposons known as starships (Urquhart et al., 2024). These mobile elements can move large stretches of DNA, often exceeding 100 kbp, which can mobilize virulence-associated genes and have been associated with large structural variations in other fungi (Bucknell et al., 2025; Gluck-Thaler et al., 2022; F. Liu et al., 2025). After performing a broader search using starfish, we were unable to confirm the insertion site of the putative starship in *A. kierstenii* sp. nov. Starfish utilizes multiple genomes to confirm the presence of these elements and is best suited to closely related genomes, hence its ability to locate starships in both *A. alternata* strains (Gluck-Thaler & Vogan, 2024b). Manual curation was able to locate a myb/SANT domain protein which often corresponds to the end of a starship and, as such, we maintain that this element represents a full, untruncated starship (O’Donnell et al., 2025).

Analysis of the cargo content detected multiple virulence-associated genes, including a tandem repeat containing multiple copies of a GMC oxidoreductase, a cytochrome p450 enzyme and a haloacid dehalogenase. GMC oxidoreductases are most often associated with lignocellulose degradation in various fungal species through the production of hydrogen peroxide (Sützl et al., 2018). The remaining two proteins have been shown to have many differing functionalities, including virulence and detoxification of various metabolites (Chen et al., 2014; D. Li et al., 2025). While the impact of this tandem duplication event on pathogenicity is unknown, the capture of these enzymes by a transposase itself also poses potential biosecurity threats. This is reminiscent of the capture and subsequent movement of toxA via a Tc1 transposon within a large starship labeled Sanctuary (Bucknell et al., 2025). This starship then shuttled *toxA* across multiple wheat-infecting species, including an *Alternaria* species (F. Liu et al., 2025). Future investigations will focus on determining the potential spread of this transposon to other *Alternaria* species and its effects on pathogenicity.

*Alternaria kierstenii* sp. nov. was isolated from a relatively limited but severely diseased garlic mustard population that represented only a small portion of a larger forest infestation. This infestation occurred in a truly non-managed system, with no nearby large-acreage *Brassica* crop fields. Follow-up studies examining the epidemiology of *A. kierstenii* sp. nov. or its movement into the larger contiguous garlic mustard infestation were not conducted, and there is currently no evidence that the pathogen spread beyond this localized outbreak. Amplicon sequencing data from a nationwide garlic mustard survey suggests that the association between garlic mustard and an *A. kierstenii*-like taxon may be geographically restricted to the eastern United States. This pattern is consistent with expectations for a recently established host-microbe association and a putative pathogen emergence event. Within that dataset, OTU 1 was the rarest *Alternaria* clade and was detected only in samples from Pennsylvania, New York, and Maryland. Of the two associated ASVs, one was a 100% sequence match to *A. kierstenii* sp. nov. There are limitations to the use of ITS sequences alone for phylogenetic and species identification; however, only three publicly available NCBI BLASTn records shared 100% similarity with this sequence. Two of these are from *A. kierstenii* sp. nov. isolates, and one is a recently deposited aerial eDNA sequence from an uncultured fungus collected using an air filter in British Columbia, Canada. Although garlic mustard is widespread across Canada, the quality metrics and read abundance associated with the aerial eDNA record are not available, limiting interpretation of this match. In the garlic mustard microbiome dataset, additional populations also contained reads assigned to the *A. kierstenii*-like ASV. However, these detections were excluded from consideration because their read counts were low, with ≤ 15 reads per sample (n = 5), and therefore fell below our predefined stringency threshold of 100 reads per sample. The *A. kierstenii*-like ASV was the dominant *Alternaria* ASV for both locations identified in Pennsylvania and New York, representing >81% of total *Alternaria* ASVs. As all these microbiome samples were collected from asymptomatic tissue, and the Maryland outbreak appeared localized, therefore the isolate described here in detail may represent an opportunistic *Alternaria* endophyte that acquired novel pathogenicity features. Future studies are needed to recover additional *Alternaria* strains from the garlic mustard populations with matching ASVs recorded in New York and Pennsylvania, in addition to garlic mustard infestations across the eastern US, which will be critical for additional genomic studies. Using genomic datasets to investigate molecular factors influencing pathogen emergence for this new species is limited due to the availability of only a few distantly related Brassicaceae-infecting *Alternaria* genomes. Ultimately, these additional studies are critical to understanding the possible origins and distribution of *A. kierstenii* sp. nov. and/or *A. kierstenii*-like taxa potentially distributed across the US, especially in New York, which ranks second among US states in cabbage production with a 2025 production value exceeding US$59 million (USDA, 2026).

While much attention has focused on the agroecological impact of invasive non-native plant species (i.e., weeds) (Pimentel et al., 2005; Vilà et al., 2011), the biosecurity risks of microbial movement and plant pathogen evolution remain overlooked for these widespread, dense weed infestations. The characterization and evaluation of *A. kierstenii* sp. nov., a newly described brassicaceous plant infecting species isolated from the invaded range of garlic mustard, underscores the agricultural risks posed by invasive weeds and the need for proactive pathogen surveillance. Compared with crop diseases, diseases of invasive or wild plant populations remain an underexplored area of phytopathology, largely because these invasive hosts are infrequently recognized as important to the health of individual commodities or other crop hosts. However, the biosecurity risks associated with pathogen spillover from invasive weed reservoirs into taxonomically related, sympatric cropping systems remain persistent and highly relevant to US agriculture. Host density plays a major role in disease ecology, and unlike naturalized weedy plants that are typically scattered within or adjacent to agricultural fields, invasive species can form widespread, dense monocultures (McDonald & Stukenbrock, 2016). Long-lived monoculture populations that harbor diverse and novel pathogens, and that can serve as ancillary hosts for crop pathogens, create disease-conducive conditions that favor pathogen selection, evolution, and emergence (Fabiszewski et al., 2010; Goss et al., 2020; Linde et al., 2016). These risks are further compounded by significant uncertainties, including the limited characterization of weed-associated microbiota that may be introduced, maintained, or dispersed across invaded regions (Fulcher et al., 2024). While still abstract in most cases, concerns about the agricultural risks posed by invasive plant species are warranted, especially when considered alongside documented wildlife spillover events that have caused severe pandemics in diverse animal systems and driven zoonotic disease outbreaks, including some of the most lethal diseases affecting human and livestock populations (Chai et al., 2023; Jori et al., 2021; Saldaña et al., 2024).

## DATA AVAILABILITY

The newly sequenced garlic mustard-isolated *A. kierstenii* sp. nov. genome assembly and annotations are made available through the USDA Ag Data Commons repository. These data are currently accessible with a temporary, private link and can be retrieved through the Figshare platform (https://figshare.com/s/5aeee14a5a4b68a7148c).

## ACKNOWLEDGEMENTS

We thank Brewster F. Kingham and Olga Shevchenko at the University of Delaware DNA Sequencing & Genotyping Center for their valued assistance. Funding for this research was supported by the United States Department of Agriculture (USDA), Agricultural Research Service (ARS) through project 8044-30400-001-000-D, Animal and Plant Health Inspection Service through project 8044-22000-047-022-I, and in part by an appointment of C. Sawyer to the ARS Research Participation Program administered by the Oak Ridge Institute for Science and Education through the interagency agreement between the U.S. Department of Energy and the USDA. ORISE is managed by ORAU under DOE contract number DE-SC0014664. Portions of this manuscript text were edited with the assistance of large-language models implemented though USAi (https://chat.usda.usai.gov). The findings and conclusions in this publication are those of the authors and should not be construed to represent any official USDA, DOE, or U.S. Government determination or policy. The mention of trade names or commercial products in this publication is solely for the purpose of providing specific information and does not imply recommendation or endorsement by the U.S. Department of Agriculture. USDA is an equal opportunity provider and employer.

## SUPPLEMENTARY TABLES

**Supplementary Table 1.** Plant species tested in this study.

| **Scientific Name** | **Common Name** | **Cultivar** | **Seed Source<sup>\*</sup>** | **Status** |
|---|---|---|---|---|
| *Alliaria petolata* | Garlic mustard | - | Field collected - Frederick, MD | Invasive |
| *Brassica oleracea var. capitata* | Cabbage | Primo Vantage | Johnny's Selected Seeds | Food crop |
| *Guillenia flavescens* | Field mustard | - | Johnny's Selected Seeds | Cover crop |
| *Solanum lycopersicum* | Tomato | Mountain Fresh Plus (F1) | Johnny's Selected Seeds | Food crop |
| *Raphanus sativus* | Radish | Crunchy King (F1) | Johnny's Selected Seeds | Food crop |
| *Matthiola incana* | Stock | Ames 26267 | NCRPIS | Ornamental crop |
| *Erysimum cheiri* | Wallflower | Ames 21403 | NCRPIS | Ornamental crop |
| *Capsicum annuum* | Bell pepper | Ace (F1) | Johnny's Selected Seeds | Food crop |

<sup>\*</sup>Sources: Johnny's Selected Seed (Winslow, ME);  North Central Regional Plant Introduction Station (NCRPIS), USDA-ARS (Ames, IA)

**Supplementary Table 2.** Starships annotated in this study.

| **#elementID** | **elementCaptainID** | **elementContigID** | **elementBegin** | **elementEnd** | **elementLength** | **elementStrand** | **emptyContigID** | **emptyBegin** | **emptyEnd** | **emptyLength** | **emptyStrand** | **emptySiteSeq** | **totalFlankAlignment** | **quality** | **warnings** |
|---|---|---|---|---|---|---|---|---|---|---|---|---|---|---|---|
| PN1_s00030 | PN1_tyr10 | PN1_scaffold2 | 3,498,061 | 3,617,940 | 119,880 | + | Z7_scaffold2 | 2,025,302 | 2,025,311 | 10 | + | tgaaaattag | 197,583 | uniq | . |
| PN1_s00031 | PN1_tyr11 | PN1_scaffold2 | 3,498,061 | 3,617,940 | 119,880 | - | Z7_scaffold2 | 2,025,302 | 2,025,311 | 10 | - | tgaaaattag | 197,583 | uniq | . |
| Z7_s00035 | Z7_tyr15 | Z7_scaffold8 | 1,735,552 | 1,823,050 | 87,499 | + | PN1_scaffold9 | 1,727,864 | 1,727,873 | 10 | - | tgaaaattag | 195,515 | uniq | . |
| Z7_s00036 | Z7_tyr16 | Z7_scaffold8 | 1,735,552 | 1,823,050 | 87,499 | - | PN1_scaffold9 | 1,727,864 | 1,727,873 | 10 | + | tgaaaattag | 195,515 | uniq | . |

## SUPPLEMENTARY FIGURE LEGENDS

**Supplementary Figure 1.**
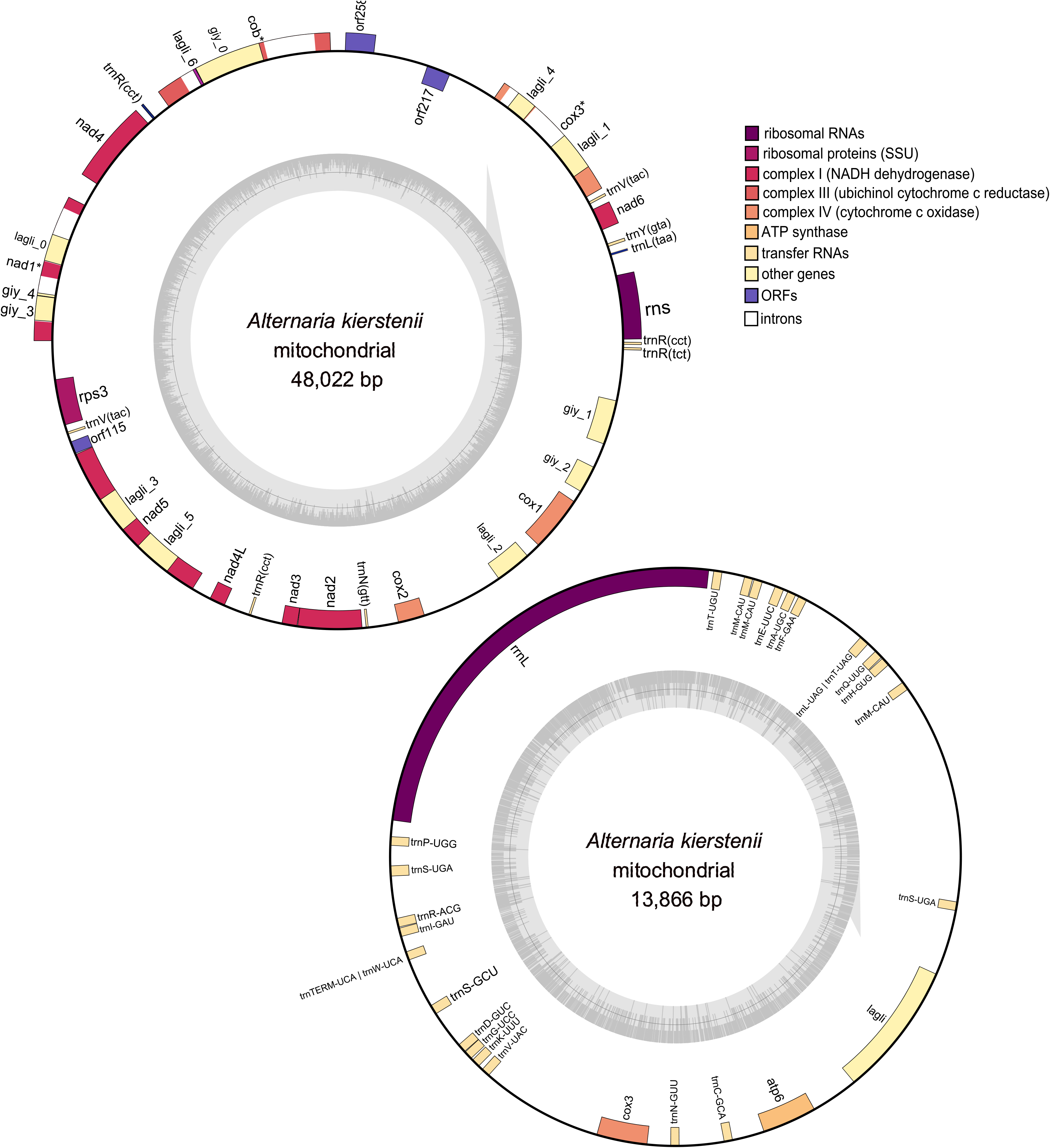
Mitogenome of *Alternaria kierstenii* sp. nov. Outer circle represents the coding sequence with gene features and tRNAs labeled. Inner circle represents the GC content of each contig.

**Supplementary Figure 2.**
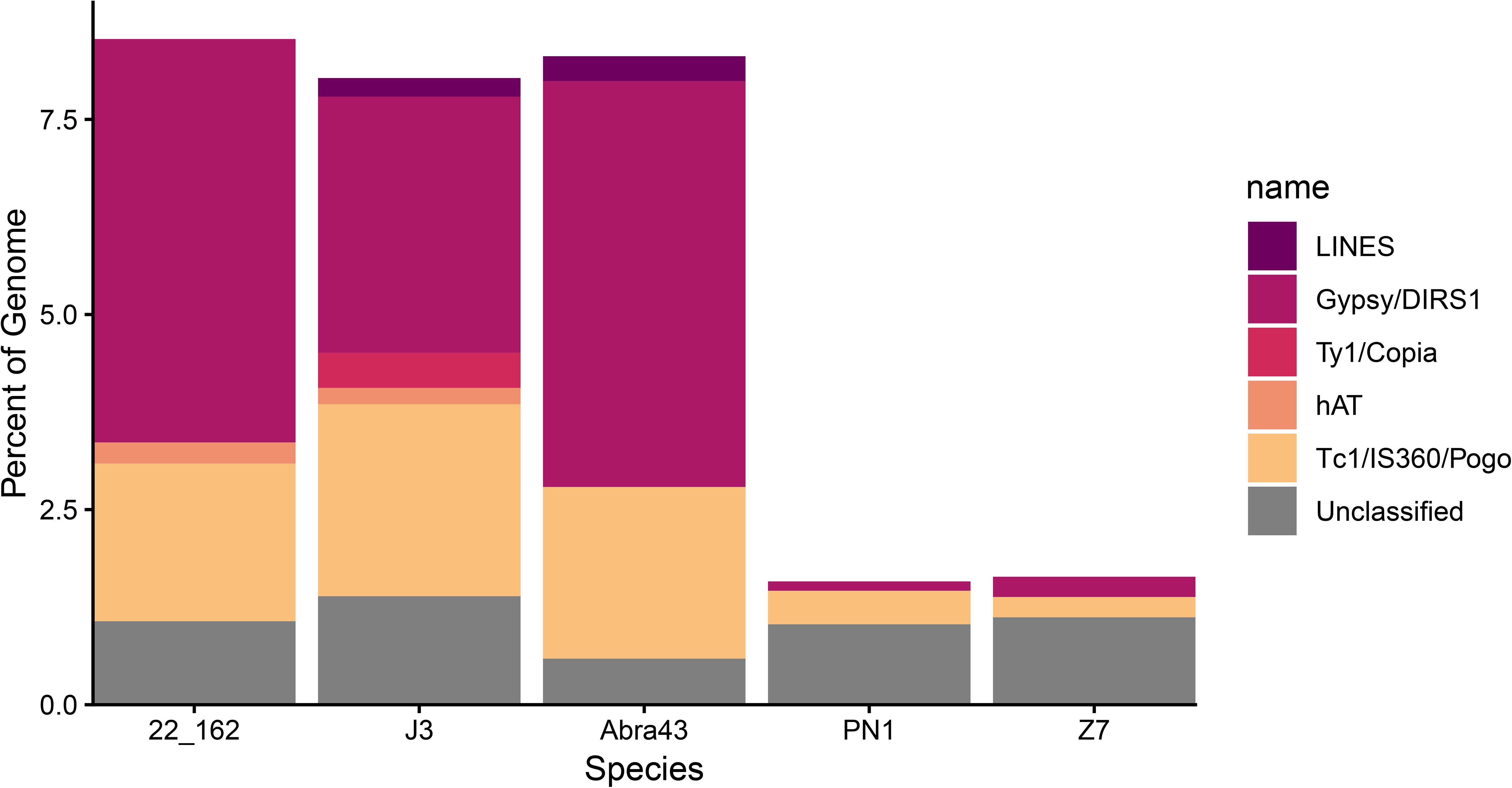
Stacked bar charts of repetitive elements across the fungal cohort as predicted with repeatmodeler. Bars represent percent of the genome composed of a given repetitive element. 22-162 – *A. kierstenii* sp. nov. FDWSRU 22-162, J3 – *A. brassicae* J3, Abra43 – *A. brassicicola* Abra43, PN1 – *A. alternata* PN1, Z7 – *A. alternata* Z7.

**Supplementary Figure 3.**
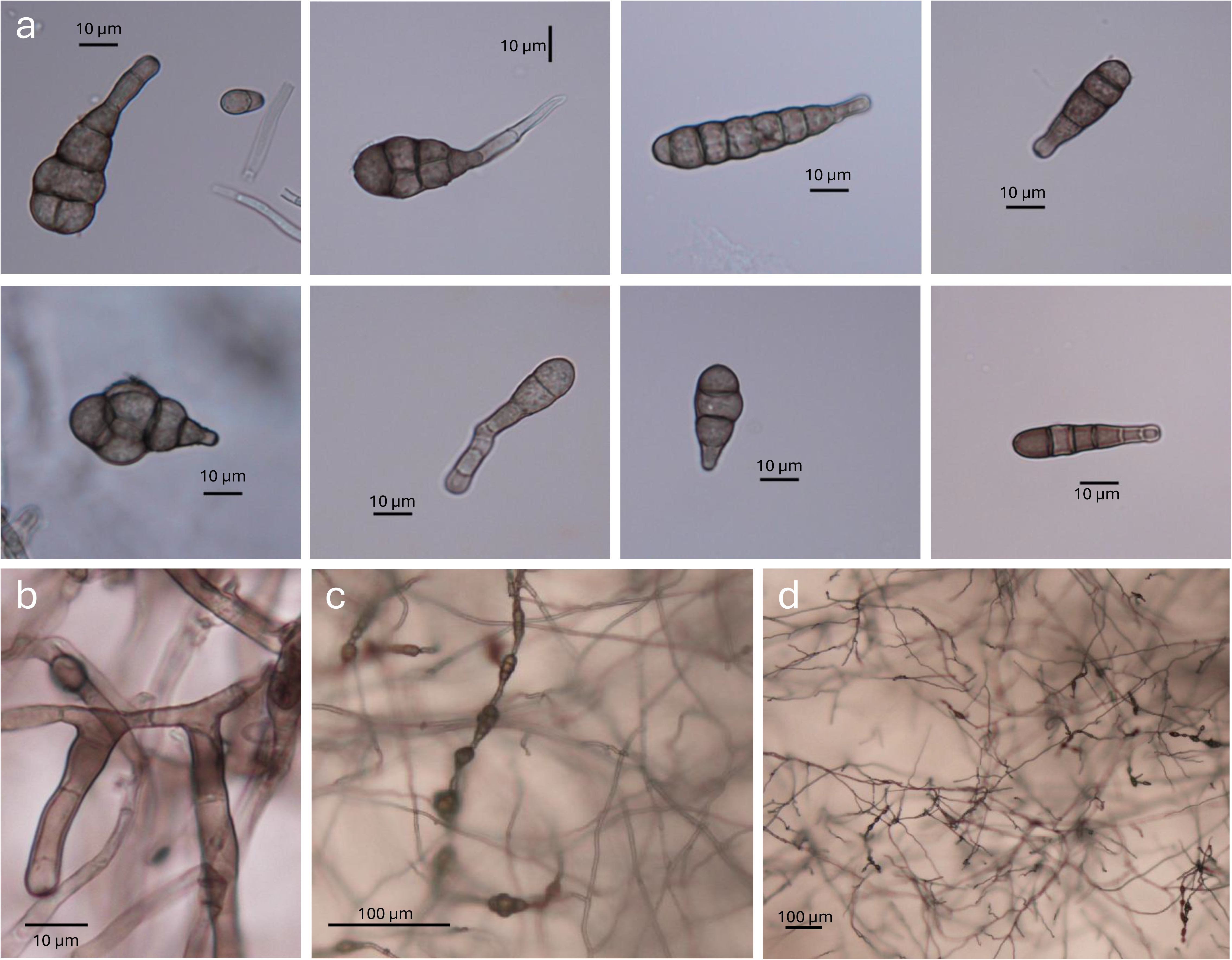
Spore phenotype of *Alternaria kierstenii* sp. nov. a) Eight representative images of conidia isolated from cultures of *A. kierstenii* sp. nov. b) Conidiophore of *A. kierstenii* sp. nov. c-d) Images of spore chains from a culture of *A. kierstenii* sp. nov.

